# Linking histology and molecular state across human tissues

**DOI:** 10.1101/2022.06.10.495669

**Authors:** Andrew Jones, Gregory W. Gundersen, Barbara E. Engelhardt

## Abstract

Histological imaging and molecular profiling of human tissues both offer information-rich characterizations of biological structure and function. Each of these modalities has been used to characterize the organization and dysregulation of a variety of tissues and cell types. While large-scale studies of each modality in isolation have been conducted, it remains largely unknown the extent to which these two views of a tissue relate to one another. Understanding how cellular states are encoded in cellular morphology would increase the utility and interpretability of imaging data; conversely, understanding the state of the cells within histology images would give deeper insights into the types and states of cells that constitute these tissue samples. To this end, we jointly analyzed 13, 360 human tissue samples with paired bulk gene expression profiles and histology images across 935 donors from the Genotype and Tissue Expression (GTEx) Consortium v8 study. This analysis reveals relationships among gene expression and cellular morphology through shared sources of expression and morphological heterogeneity both within and between tissue types. We describe shared sources of variation including cell-type heterogeneity, sample ischemic time, and donor health and demographics. We find specific correlated effects in both morphology and transcription linked to specific donor characteristics, such as their use of mechanical ventilation. This paired understanding adds value to each data modality on their own by enabling a more precise characterization of the alternative modality in the absence of those data.

## 1 Introduction

Histology images have been used for many purposes in the biological and medical sciences, from studying the basic biology of cellular morphology in the lab (Nedzved et al., 2000; Phoulady et al., 2016; Sornapudi et al., 2018) to identifying and diagnosing cancerous tumors in the clinic (Cireşan et al., 2013; Araújo et al., 2017; Aresta et al., 2019). Recently, computational approaches have become essential tools for analyzing high-resolution histology images (Fuchs and Buhmann, 2011; Komura and Ishikawa, 2018; Gratz et al., 2020; Barisoni et al., 2020). While these approaches initially relied on hand-crafted image features, there has been a recent push to apply modern machine learning methods that automatically extract salient features of the data (Van Valen et al., 2016; Moen et al., 2019).

In 2012, the computational vision community developed representations of images estimated using deep neural networks (DNNs) that outperformed representations based on hand-crafted features on important downstream tasks, including classification, prediction, and clustering for community benchmarks (Krizhevsky et al., 2012). A major downside of DNN-based models, which often have thousands or millions of parameters, is the need for large sample sizes in order to perform inference. Moreover, with a few notable exceptions (Gundersen et al., 2019; Ash et al., 2021; Fu et al., 2020), the methods developed for histology image analysis have focused on prediction and classification rather than unsupervised characterization of the variation among the images or image features (Cireşan et al., 2013; Araújo et al., 2017; Aresta et al., 2019; Fuchs and Buhmann, 2011).

Advances in machine learning and computer vision (Krizhevsky et al., 2017; He et al., 2016; Kingma and Ba, 2014), along with the development of large-scale histology and pathology image collections (GTEx Consortium et al., 2017, 2020; Network et al., 2012), have catalyzed huge strides in automated classification and diagnosis of diseased tissues by applying machine learning to histology images. For example, machine learning systems can leverage huge databases of histology images to classify tissues as tumorous or benign, identify dermatological diseases, and segment and identify cell types in tissues (Beck et al., 2011; Bejnordi et al., 2017; Kothari et al., 2013). In parallel, the rise of large-scale molecular profiling of tissue samples has provided a distinct and complementary window into the biological processes of healthy and diseased tissues (GTEx Consortium et al., 2017, 2020). Gene expression studies in particular have provided valuable insights into cellular function in both healthy and diseased tissues (GTEx Consortium et al., 2020; ICGC/TCGA Pan-Cancer Analysis of Whole Genomes Consortium et al., 2020). However, it remains largely unknown how these molecular-level features relate to the morphology of a tissue sample. Understanding this relationship could improve disease diagnosis and suggest more effective interventions using only inspection of these images. Furthermore, a characterization of this relationship could also reduce the need for molecular profiling when histology images contain signatures of the relevant molecular markers. Overall, it would be beneficial to treat these modalities as complementary to one another — understanding what information is jointly encoded in both and what information is exclusively encoded in one or the other. This joint understanding could be a crucial step toward piecing together a comprehensive picture of the state of a tissue sample.

Several lines of work have used dimension reduction techniques to study the associations between genomic measurements and cellular morphology. A handful of studies have specifically focused on studying joint variation in cancer cells. One approach leveraged data from The Cancer Genome Atlas (TCGA) to identify clusters of glioblastoma subtypes that showed distinct morphology characteristics and molecular features (Cooper et al., 2012). Another effort studied the relationship between histology images and expression in TCGA, identifying correlation between image features and expression of a set of genes that are known to be linked to cancer patient outcomes (Subramanian et al., 2018). Another line of work identified imaging biomarkers associated with immune cell infiltration in thyroid tissue from healthy patients (Barry et al., 2018).

Two recent studies used variations of canonical correlation analysis (CCA) (Hotelling, 1992) to examine the associations between bulk gene expression and histology image features in GTEx v6 data. The first, which is called ImageCCA, identifies joint variation between gene expression profiles and image features (Ash et al., 2021). To generate image features, ImageCCA applied a convolutional autoencoder (CAE) to produce a low-dimensional embedding for each image. Examining the factors estimated using ImageCCA, the authors found several shared patterns between the data modalities related to tissue type. Furthermore, the ImageCCA analysis identified image morphology QTLs (imQTLs), or associations between donors’ genotypes and paired image features, in the GTEx v6 data. The authors also applied their method to data from two separate studies in The Cancer Genome Atlas (TCGA) (Weinstein et al., 2013). A gene set enrichment analysis on the estimated factors revealed substantial covariation between expression and images features associated with extracellular processes and cell-type heterogeneity in TCGA.

Building off of ImageCCA, a second CCA-based approach was developed that simultaneously extracts salient image features and identifies their association with gene expression profiles (Gundersen et al., 2019). This method — called deep probabilistic canonical correlation analysis (DPCCA) — jointly fits an image autoencoder and CCA model. Applying DPCCA to the GTEx v6 data, the authors found that many of the latent factors were driven by differences between tissue types, in contrast to results using the two-stage approach of ImageCCA. A QTL analysis using the DPCCA latent variables as phenotypes revealed several imQTLs. While both of these studies reveal important links between gene expression, image morphology, and genotypes, they rely on the limited GTEx v6 data. As convolutional autoencoders are known to require large sample sizes to achieve reasonable out-of-sample performance because of the large numbers of parameters, we are interested in applying these approaches to much larger paired imaging and gene expression datasets.

To do this, we consider the GTEx v8 data, which contain six times the number of paired gene expression and histology image samples compared to GTEx v6 (GTEx Consortium et al., 2020). The GTEx v8 data contain 13, 360 paired expression profiling and histology samples in 55 tissues from 935 donors. The unprecedented size of this multi-modal dataset offers an opportunity to study how molecular characteristics of tissue samples relate to the complex morphology observed in images of those same tissues. A joint “multi-view” analysis of these two modalities makes it possible to understand each view better, as well as to provide insights into the system that are difficult or impossible to identify when using only a single view. Thus, we aim to analyze shared biological variation in the GTEx v8 sample with paired histology images and bulk gene expression profiling; we also leverage the available genotype data for each donor.

This paper proceeds as follows. First, we present a joint analysis of the GTEx v8 histol-ogy images and expression profiles using latent variable models that uncover the sources of shared variation between the two paired data modalities. We then examine the biological origins of some of the most dominant sources of variation, including cell-type heterogeneity and tissue type. We also analyze technical variation in the data modalities, including ischemic time and patient ventilator status, and how these covariates affect the relationship between morphology and expression. Leveraging the genotype data, we find a number of quantitative trait loci that describe associations between certain genotype variants and image morphology. Using modality imputation methods, we show that our approaches are able to reveal biological insights using one data modality that were previously available only through access to the other data modality.

## 2 Results

We examined the relationship between tissue histology and molecular features using the GTEx v8 data. To this end, we fit dimension reduction methods on the paired histology image and bulk gene expression samples, and we used the lower-dimensional representations of these methods to explore different sources of variation: i) which gene expression patterns are associated with tissue morphology patterns; ii) whether there are any genetic variants that are jointly associated with tissue histology and gene expression; and iii) how biological and technical covariates affect the shared variation of the two data modalities.

### 2.1 The GTEx v8 data include joint observations of genotype, bulk gene expression, and histology images across diverse healthy human tissues

The GTEx v8 dataset contains paired measurements on 13,360 samples across three modalities: histology images, RNA-seq bulk gene expression profiles, and genotypes. The GTEx v8 data, which come from 935 donors, span 55 distinct healthy human tissues (Figure 1). The v8 data contain over six times as many paired observation samples as the previous GTEx v6 data, which had 2221 paired samples across 499 individuals.

**Figure 1:**
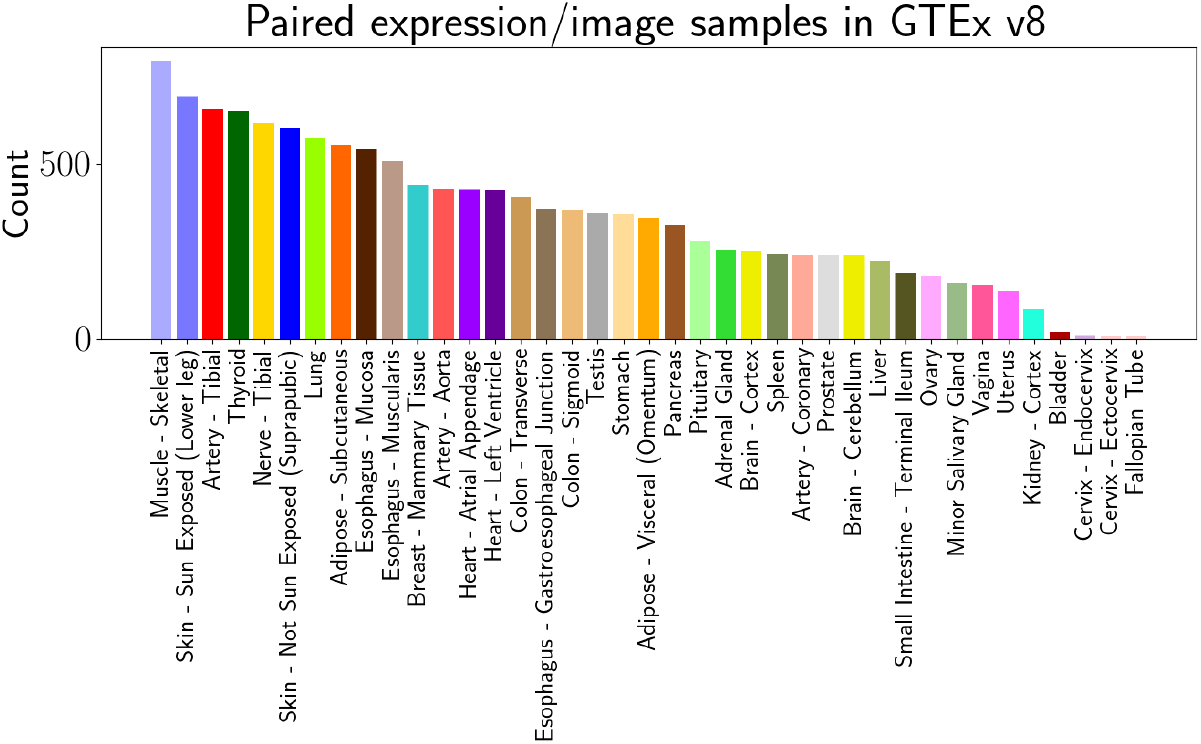
Makeup of the GTEx v8 dataset. (a) Counts of the number of samples and tissue types with paired measurements of image morphology and gene expression. The GTEx v8 dataset contains 13,360 samples with joint data modalities, which is over six times as many samples as the GTEx v6 data. (b) Number of paired samples for each tissue type.

The imaging data were collected from tissue slices fixed to slides and stained with hematoxylin and eosin (H&E). Raw images used inconsistent magnification, but in general were taken with either 20× apparent magnification, resulting in around 0.25 microns per pixel, or 40× apparent magnification, resulting in around 0.50 microns per pixel. Each image was post-processed to have a size of 1000 × 1000 pixels. These imaging data exist to ensure that each of the RNA-sequencing samples was from healthy tissue (Carithers et al., 2015), as the goal of GTEx was to survey and sequence non-diseased tissues from postmortem donors. Only solid tissue samples were imaged, restricting our paired sample data to 39 tissues. The tissue slices for imaging and for bulk RNA-sequencing were harvested from adjacent locations in the tissue of interest. For our analyses, we assume that these paired slices to represent the same section of tissue.

The bulk RNA-sequencing data were measured in transcripts-per-million (TPM). As in earlier work, we log-transformed and z-scored the RNA-seq data for our analyses (Ash et al., 2021). The genotype data was based genotype calls from OMNI SNP arrays. Variants with a minor allele frequency (MAF) of ≤ 1% were excluded. The donor demographic information and medical history were ascertained using postmortem surveys of next-of-kin (Carithers et al., 2015).

### 2.2 Separability of tissue types

We first sought to understand the major sources of variation within the imaging and gene expression data. To do this, we fit dimension-reduction methods to each: for the imaging data, we fit a convolutional autoencoder (CAE) (Masci et al., 2011) followed by dimension reduction using UMAP (McInnes et al., 2018) on the CAE embeddings; for the gene expression data, we fit UMAP directly on the log-transformed and z-scored gene expression values (Figure 2). We found that tissue type is the driving factor of variation in the gene expression data. The UMAP embedding of the expression data easily separated the samples by the tissue types (Figure 2a). Meanwhile, the embedding for the image data did not show a clear division among the tissue types (Figure 2b). Some tissues, such as testis and mammary gland, showed strong within-tissue coherence in the projection of the images, but most tissue types did not show strong within-tissue clustering. However, examining rela-tionships across tissues, we observed that some tissues with similar visual appearances — such as skeletal muscle, heart, and thyroid — were well-mixed in the embedding. For both data modalities, we computed the adjusted Rand index (ARI) for a K-means clustering to quantify the clustering of tissues in the two-dimensional UMAP embedding (Figure 2c); we find that tissues cluster well in the gene expression data (ARI 0.64) whereas there is poor tissue clustering in the image feature projection (ARI 0.19).

**Figure 2:**
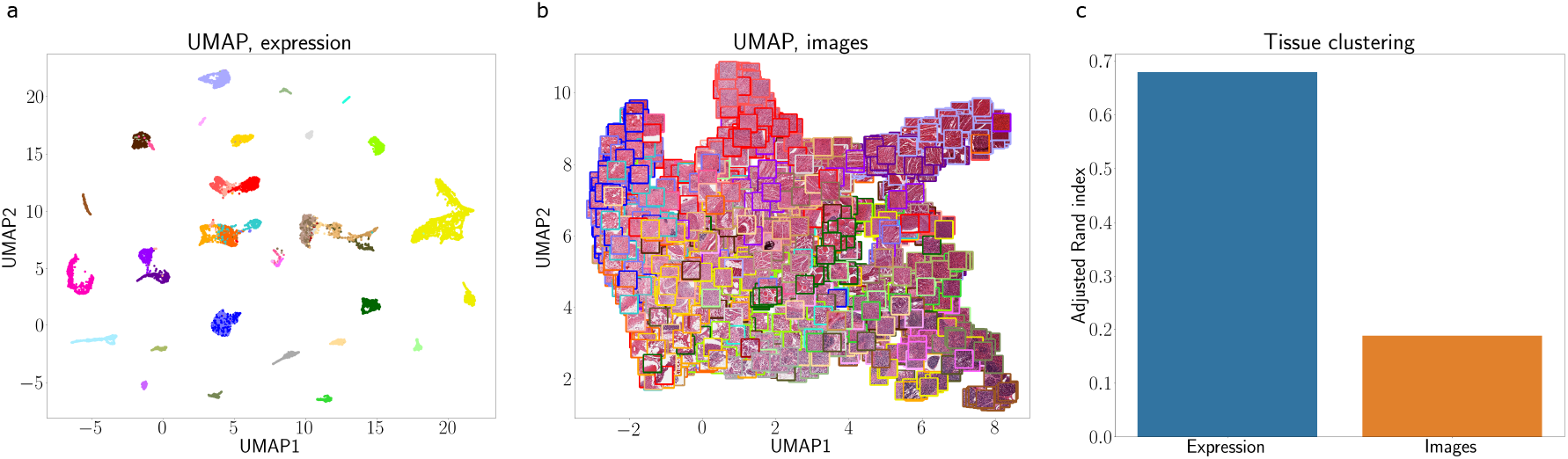
Low-dimensional embeddings of gene expression and histology image data. (a) Two-dimensional UMAP embedding of the gene expression profiles. Points (samples) are colored by their tissue types (colors correspond to those shown in Figure 1). (b) Two-dimensional UMAP embedding of the convolutional autoencoder histology image representation. The border of each image is colored according to its tissue type. (c) Adjusted Rand index computed on a K-means clustering of each UMAP embedding.

To further quantify the extent to which tissue type is encoded in the imaging and gene expression data, we fit a multi-layer perceptron (MLP) classifier to each modality, using the gene expression profiles and autoencoder embeddings as input. We then measured their performance on a set of held-out samples. We found that the gene expression tissue classifier performed well, with an average accuracy of 97%, while the classifier for the images achieved 38% accuracy (Figure 3a). Moreover, the confusion matrix for the expression data showed little error (Figure 3b). In contrast, the confusion matrices for the classifier revealed several commonly misclassified tissues in the image data (Figure 3c). For example, the two types of skin tissue were frequently misclassified as one another. Together, these results suggest that gene expression data includes variation associated with tissue type, while the morphological features from our CAE embedding broadly fail to distinguish tissue types.

**Figure 3:**
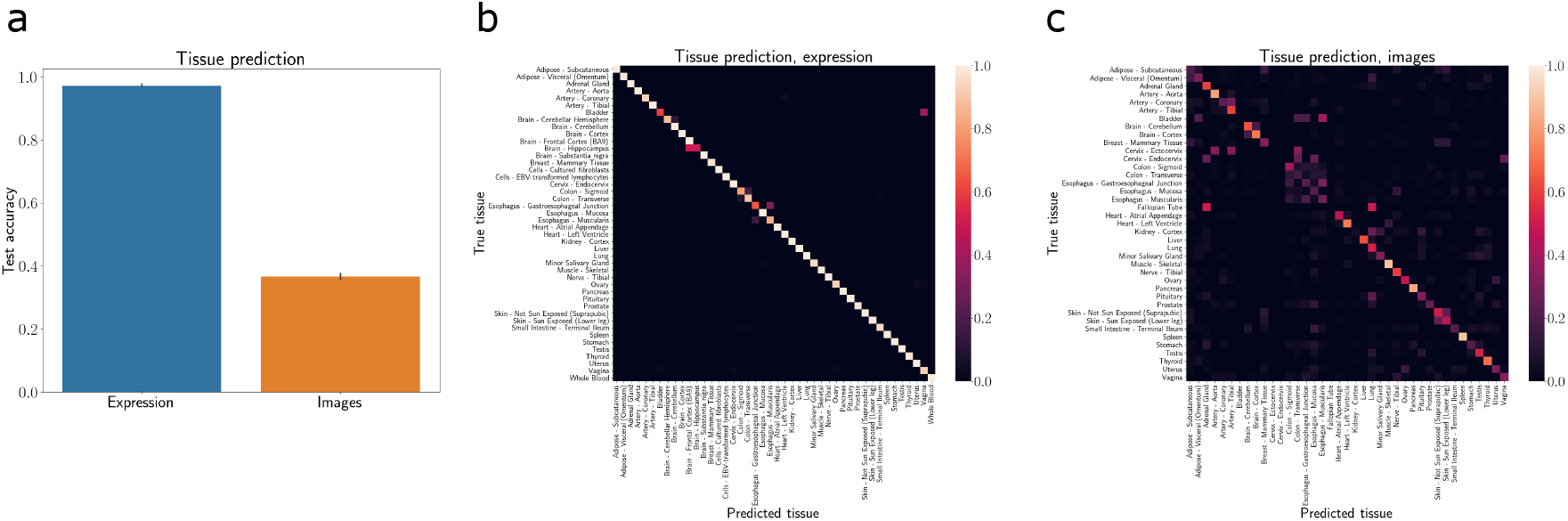
Predicting tissue type using data from the two modalities. (a) Test accuracy of a model predicting tissue type from gene expression profiles (left bar) and histology image features (right bar). For histology images, we use the autoencoder embeddings as the input to the classifier. Vertical ticks show the 95% confidence intervals for ten-fold crossvalidation. (b) Confusion matrix for predicting tissue type from gene expression profiles. The cell in row *i* and column *j* shows the fraction of samples whose true tissue type is tissue *i* that were predicted as tissue *j*. (c) Same as (b), but for predictions using autoencoder embeddings of histology images.

**Figure 4:**
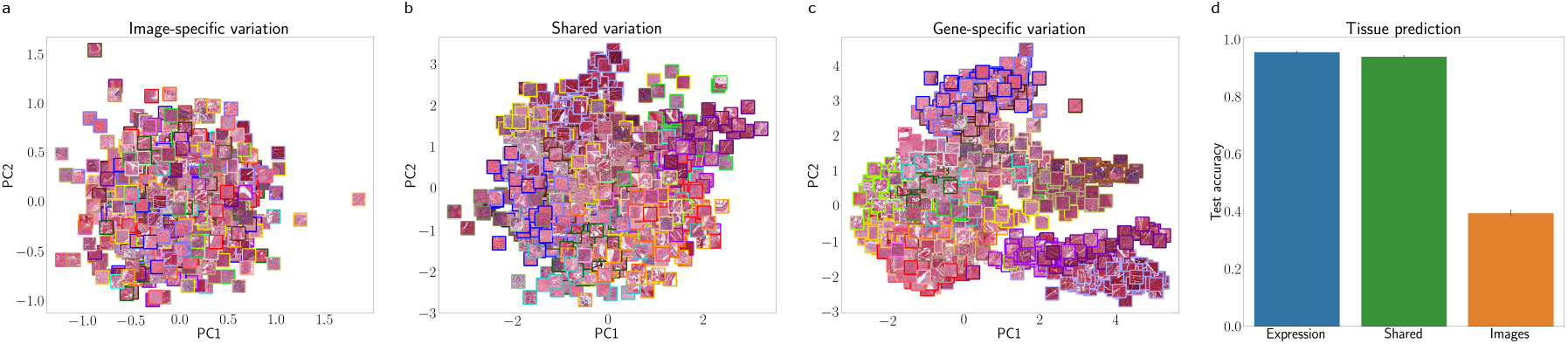
Shared variation between gene expression and histology image features captured by an IBFA model. Low-dimensional representation of each of the IBFA model’s latent variable types, capturing (a) image-specific variation, (b) variation shared between the modalities, and (c) gene expression-specific variation. (d) Test set accuracy of a model predicting tissue type from each of these latent variables. Vertical ticks show 95% confidence intervals.

### 2.3 Joint analysis of gene expression and image data

Next, we sought to explore the patterns of variation that are shared between the image and gene expression data modalities. To do this, we fit probabilistic inter-battery factor analysis (IBFA), which identifies axes of variation that are shared between two data modalities, as well as variation that is unique to each data modality. We fit IBFA using the autoencoder’s image features and gene expression profiles. The IBFA model provides a lower-dimensional representation (here, we use 50 dimensions to approximately follow previous work (Ash et al., 2021; Gundersen et al., 2019)) of each sample that takes into account the variation present in each modality.

Examining the IBFA shared latent variables, we found that many of them were driven by differences between tissue types. For example, latent dimension 21 appeared to be largely driven by ovary tissue samples (Figure 5a). Furthermore, we examined whether variation in the expression of certain genes contributed more to this latent variable than others. Using a gene set enrichment analysis (GSEA), we found an enrichment of genes with substantial contribution to this factor involved in *estrogen response*, including *AGR2* and *TFF3* (Figure 5b) (Salmans et al., 2013; Hoellen et al., 2016). These results suggest that there are substantial shared signals between the two data modalities, echoing prior work (Gundersen et al., 2019).

**Figure 5:**
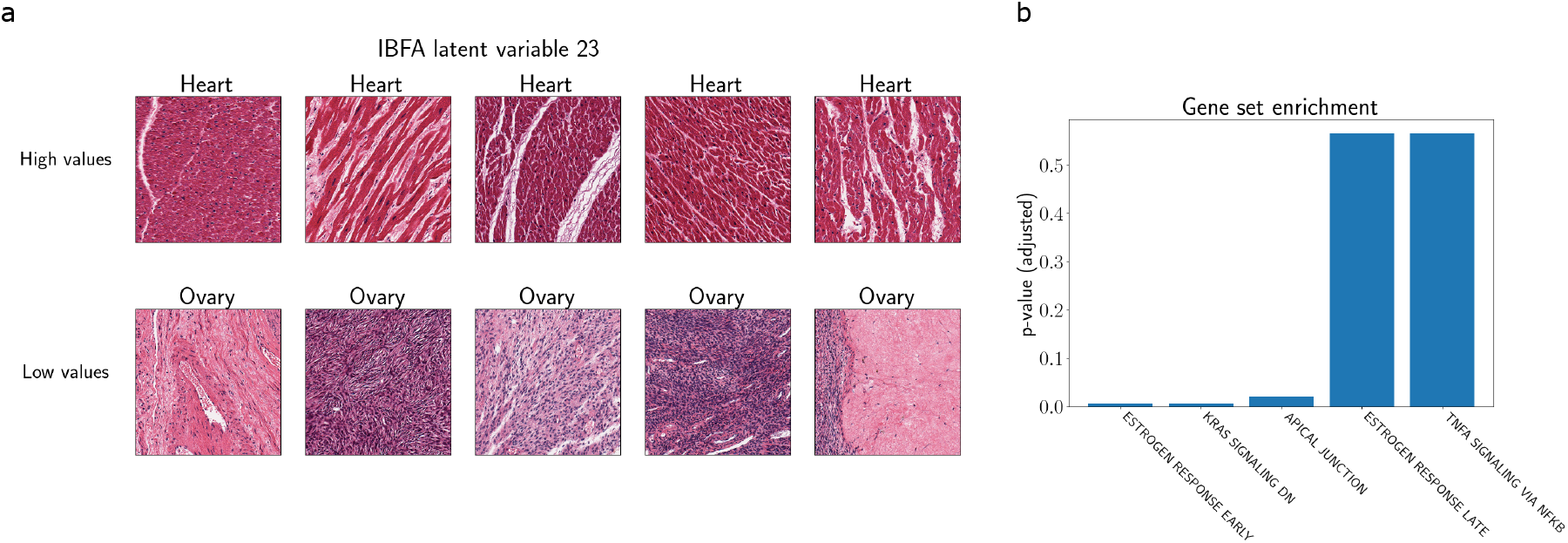
Tissue-type separation within the IBFA shared latent variables. (a) For one IBFA component (latent variable 23), the top row shows the five images with the most extreme positive estimated values for this component, and the bottom row shows the five images with the most extreme negative values. This component is likely related to heart and ovary tissue. (b) Top Hallmark pathways from a gene set enrichment analysis (GSEA) on the genes with nonzero loadings for this component. The top pathway is related to *estrogen response*.

We then examined tissue type separation in the three types of IBFA latent variables: shared, image-specific, and gene expression-specific. To do this, we fit a multilayer perceptron (MLP) classifier for each set of latent variables using the tissue type as the response, and we computed the predictive test accuracy for a set of held-out samples. We found that tissue type was well predicted using the shared latent variables and expression-specific latent variables, while accuracy was much lower for the image-specific latent variables (Figure 4d). Interestingly, prediction accuracy on held-out samples was higher for the image-specific IBFA latent variables than the prediction accuracy using the complete set of image features, as presented in the previous section. This suggests that a joint analysis of the shared and modality-specific variation between the two modalities might capture more biological signal than analyzing each data modality alone.

### 2.4 Joint associations with GTEx donor and sample covariates

In addition to exploring the relationship between histology image features and gene expression, there is a need to understand how this relationship varies across donors with varying medical health characteristics and samples with varying technical quality. To assess the paired datasets’ associations with observation covariates, we leveraged the set of donor- and sample-level metadata collected as part of the GTEx project. The donor-level metadata include characteristics such as donor demographics and medical histories, and the sample-level metadata include attributes related to sample quality and molecular information. For each metadata variable, we fit a multivariate linear regression model using the IBFA shared latent variables as covariates and the metadata variable as the response. We then measured the relationship between each latent variable and the metadata covariate using *R*^2^ of each regression model.

The most highly-associated donor-level covariates were demographic variables, such as sex (Figure 6a). Other associations were related to a donor’s overall tissue recovery and tissue health, such as ischemic time. At the donor level, ischemic time measures the interval length between between actual death, presumed death, or cross clamp application, and the start of the GTEx collection procedure. For the sample-level metadata, variables related to technical sample characteristics — such as the *rRNA rate*, or the fraction of all reads aligned to ribosomal RNA regions (Conesa et al., 2016) and the *RNA integrity number* (Schroeder et al., 2006) — showed the strongest associations (Figure 6b).

**Figure 6:**
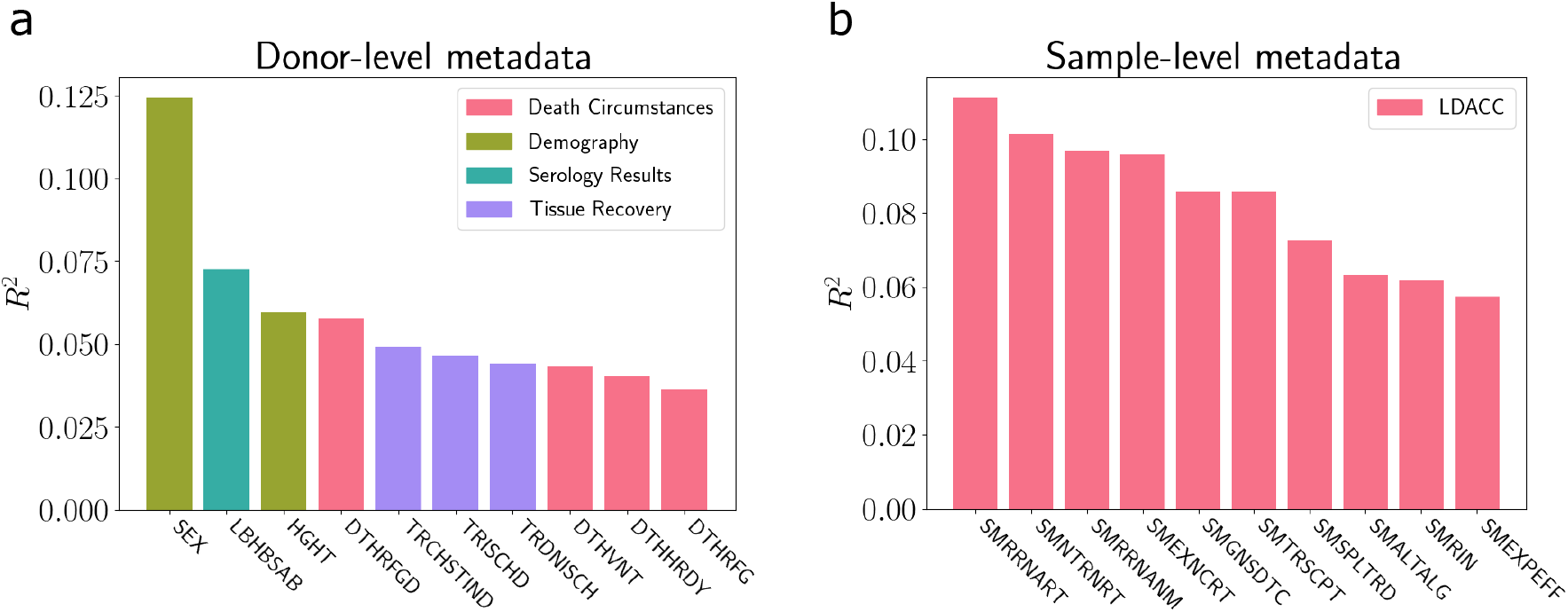
Associations between the IBFA latent variables shared between gene expression and image features, and GTEx metadata. (a) Associations between shared latent variables and donor-level metadata. Each bar shows the *R*^2^ value from a regression model fit using the IBFA shared latent variables as the covariates and the corresponding metadata variable as the response. Bars are colored by the category of metadata variable. (b) Same as (a), but for sample-level metadata. Variable descriptions are in section 5.

#### 2.4.1 Ischemic time

Motivated by the associations with ischemic time in our exploratory analysis, we began a more focused study of the association between the shared latent variables and ischemic time. Rather than quantify the association with ischemic time across all tissues as before, here, we examined each tissue’s marginal association with ischemic time. We found that the association strength varied by tissue, although most tissues showed substantial associations with ischemic time (Figure 7a). Focusing on one tissue in particular — heart tissue from the left ventricle — we found that ischemic time was most associated with latent dimension 35 (Figure 7b). Visualizing the images with values on both extremes of this factor showed visible differences between the healthy and necrotic samples (Figure 7c). These results suggest that RNA decay and tissue necrosis are jointly encoded in expression and image shared latent factors, and that those shared decay signals align well with sample ischemic time across tissues (Carithers et al., 2015).

**Figure 7:**
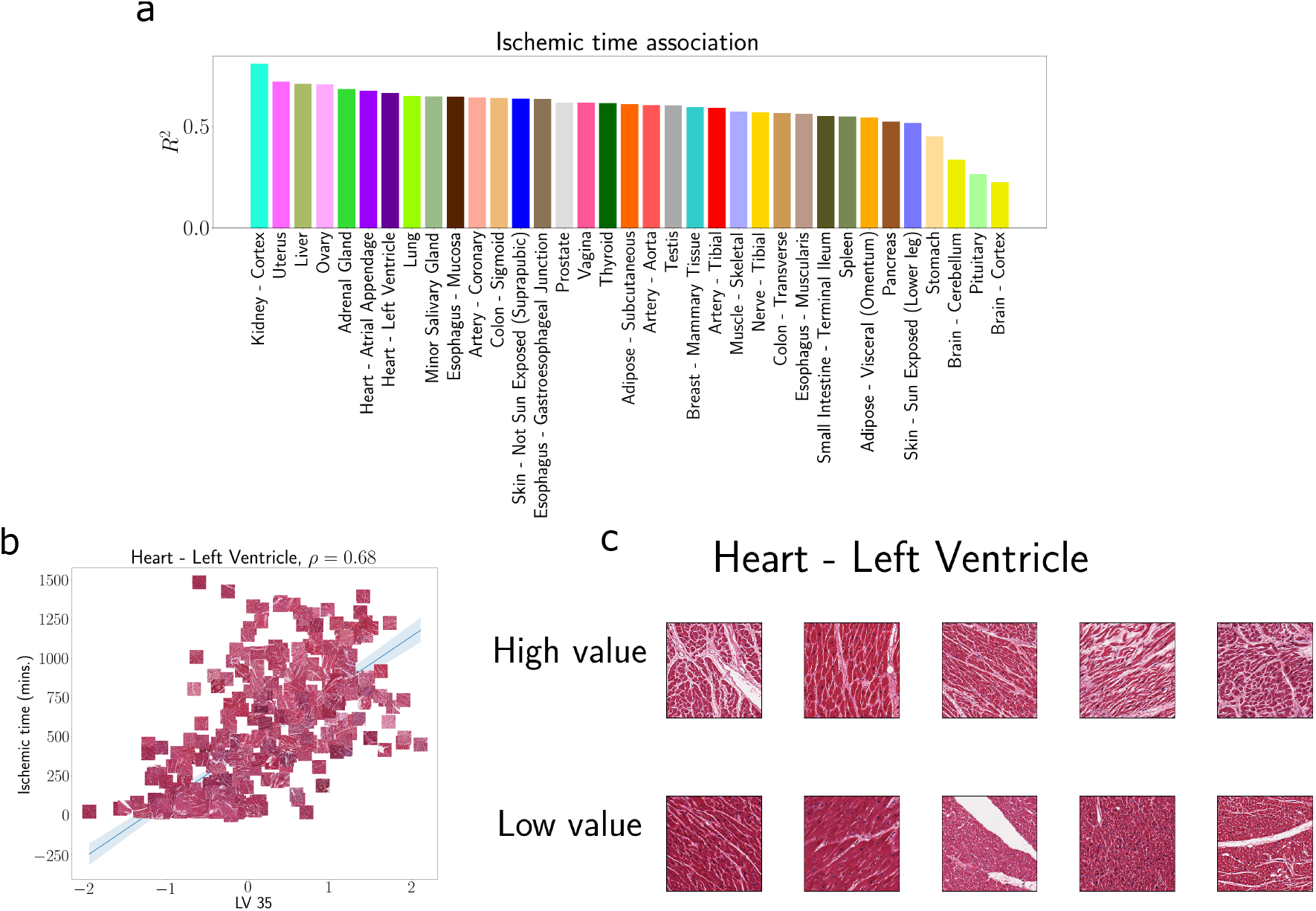
Association between sample ischemic time and shared IBFA component for samples from the left ventricle. (a) Scatterplot of one IBFA joint latent variable (LV 35) and each sample’s ischemic time for samples from the heart’s left ventricle. Each point is plotted as the corresponding histology images. (b) Histology images of the samples with the highest values (top row) and lowest values (bottom row) on this IBFA shared component.

#### 2.4.2 Donor ventilator status signal in shared variation

We next sought to explore the association between gene expression, tissue morphology, and variables related to patient medical history. In particular, we examined whether donors who were on a ventilator at the time of death showed different morphological and gene expression characteristics compared to donors who were not on a ventilator. To do so, we quantified the association between each IBFA shared component and the donors’ binary ventilator status. More precisely, for each tissue type, we fit a logistic regression model with the shared components as covariates and the ventilator status as the outcome.

We found that one shared component (latent variable 38) showed a substantial difference between these two groups of donors in lung tissue (Figure 8a). Furthermore, examining the contribution of each gene’s expression to this component, we found enrichment of genes related to *hypoxia* and *oxidative phosphorylation* (Figure 8c). One of the top genes in this component, *AQP2*, has been observed to be disturbed following mechanical ventilation (Sun et al., 2017). Another gene, *ABHD2*, was observed to correlate with risk for chronic obstructive pulmonary disease (COPD) (Liu et al., 2015). Moreover, we observed that the images of lung tissue from patients with ventilation showed different coloration and organizational patterns compared to the donors without ventilation (Figure 8b). These results suggest specific, reproducible damage done to lungs as a result of ventilation; moreover, the signal associated with ventilation shows up in both gene expression and imaging data in lung tissues.

**Figure 8:**
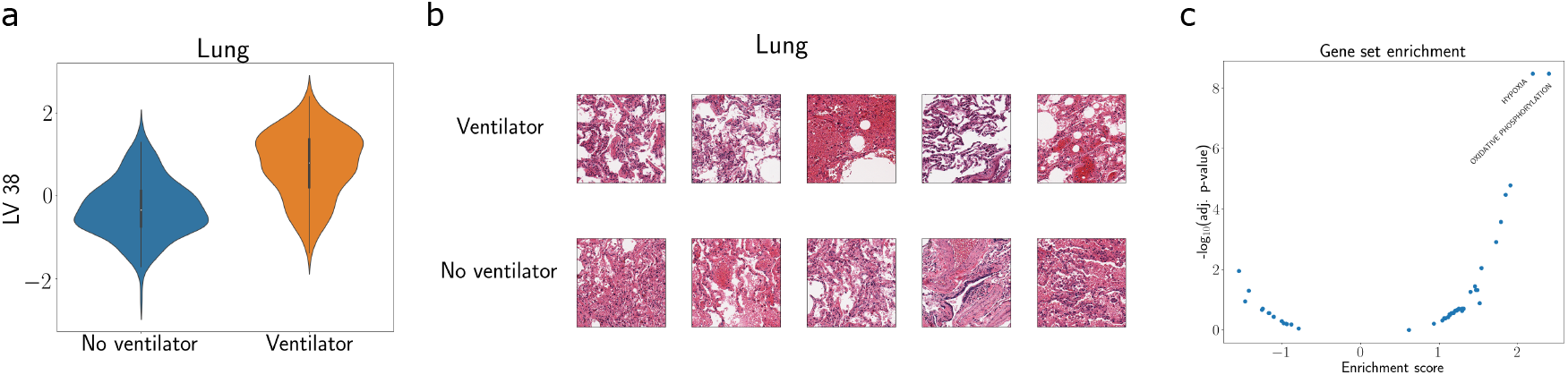
Shared IBFA component captures donor ventilator status. (a) Shared IBFA component value (component 38) for lung tissue samples from donors who were on a ventilator at the time of death (right) and those who were not (left). (b) Top row shows lung histology images from donors who were on a ventilator, with the five highest values for this component. Bottom row shows lung images from donors who were not on a ventilator, with the five lowest values for this component. (c) Gene set enrichment analysis on the set of genes with nonzero loadings for this component. The top Hallmark pathways are *hypoxia* and *oxidative phosphorylation*.

### 2.5 Image morphology QTL analysis: Incorporating genotype data

In addition to analyzing the shared morphological and transcriptional signals of these samples, we also examined these phenotypes’ associations with each donor’s genotype. Quantitative trait loci (QTL) are regulatory associations between a genetic variant and a quantative trait; the focus of GTEx has been the identification of expression QTLs (eQTLs) and splicing QTLs (sQTLs) across healthy tissues (GTEx Consortium et al., 2017). QTLs have provided valuable information about gene function and regulation (GTEx Consortium et al., 2017, 2020; Rockman and Kruglyak, 2006), in particular in the analysis and interpretation of the cellular mechanisms of genetic variants associated with disease (Cookson et al., 2009; Albert and Kruglyak, 2015). However, associations between genotype and morphological features of human tissues (image morphology QTLs, or imQTLs) have been far less studied, with a few notable exceptions (Ash et al., 2021; Gundersen et al., 2019; Barry et al., 2018).

To investigate these relationships, we conducted an imQTL analysis to discover associations between the donors’ genotypes and histology image features. For each tissue, we selected a subset of image features and genetic variants to test for association using sparse CCA and known eQTLs from GTEx (Methods). Then, we tested for association between each variant and image feature in each tissue using a linear regression model as implemented in the MatrixEQTL R package (Shabalin, 2012). We found 68 imQTL associations across six tissues (FDR ≤ 0.1), suggesting we are indeed able to use these GTEx v8 data to identify imQTLs.

As an example, the imQTLs included an association between image feature 1007 and genetic variant *rs10008860* in esophageal mucosa tissue samples (*FDR* < 0.1; Figure 9). Examining the esophageal mucosa histology images for each of these genotypes did not immediately reveal any visual differences between them; however, a more thorough analysis by a trained pathologist could be beneficial. This variant regulates the gene *AFAP1* and the long noncoding RNA *AFAP1-AS1*, which are expressed in a handful of tissues, including esophagus. Moreover, these two genes have been observed to be associated with abnormalities and disease in esophageal tissue (Wu et al., 2013).

**Figure 9:**
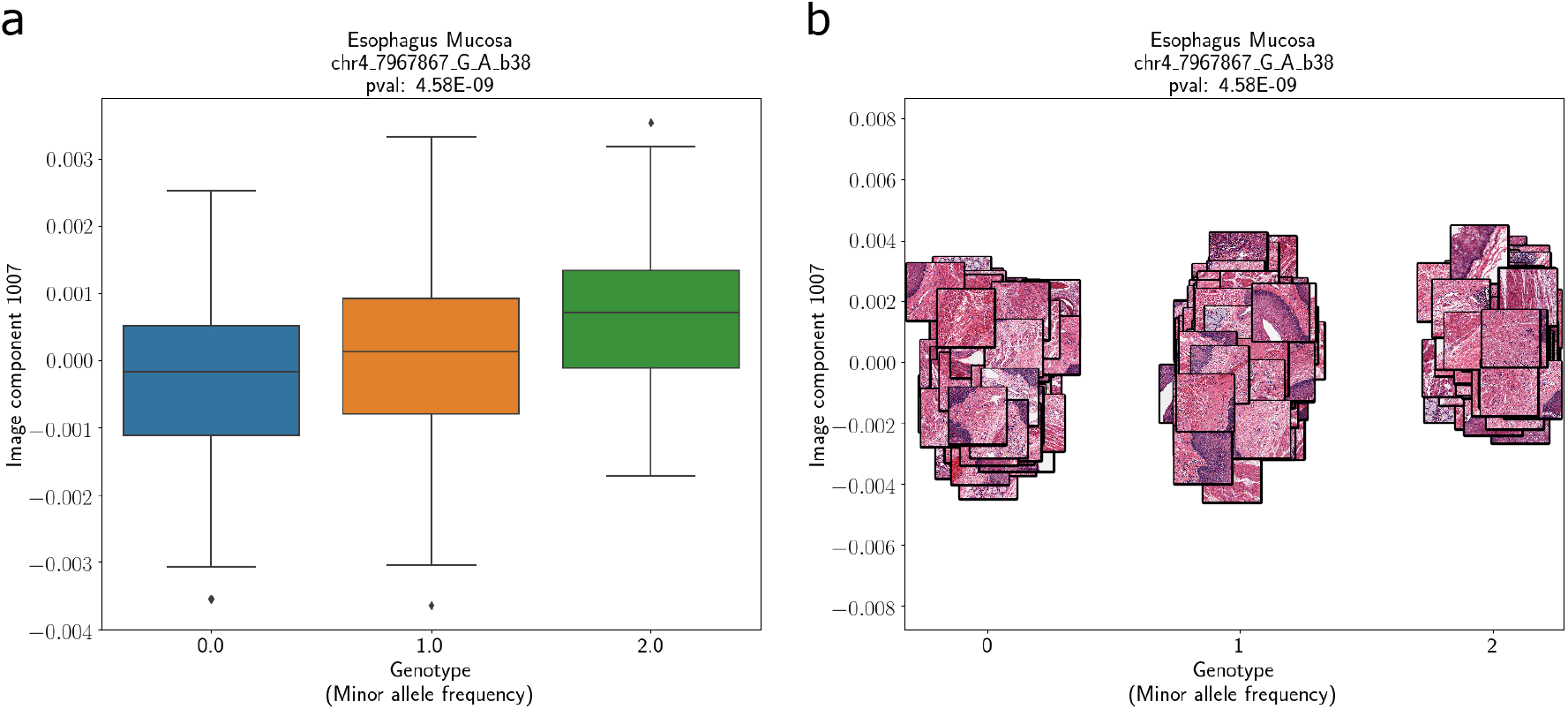
imQTL between morphology image feature and genetic variant rs10008860 in esophageal mucosa tissue. (a) boxplot with the number of minor alleles (x-axis) versus the image feature expression value (y-axis); (b) same figure but with thumbnails of the images for each sample, with jitter.

### 2.6 Sparse and dense shared components

To further explore the relationship between the image and expression datasets, we applied a matrix decomposition method that provides a mix of both sparse and dense factors, called SFAmix (Gao et al., 2013). Since SFAmix decomposes a single matrix — unlike IBFA, which jointly models two data matrices — we concatenated the expression matrix and the image autoencoder embeddings according to their matching samples. We normalized each dataset separately and fit the model with 1000 components.

Examining the resulting latent factors, we find that the majority of the factors (97%) are found to be sparse. Moreover, we examined the level of sparsity for each feature in the loadings matrix, and we found that loadings vectors corresponding to histology images tend to be much more sparse than those for gene expression (Figure 10b). This observation suggests that the shared variation between these two modalities is largely driven by many genes’ expression levels, but tends to be specific to certain image features.

**Figure 10.**
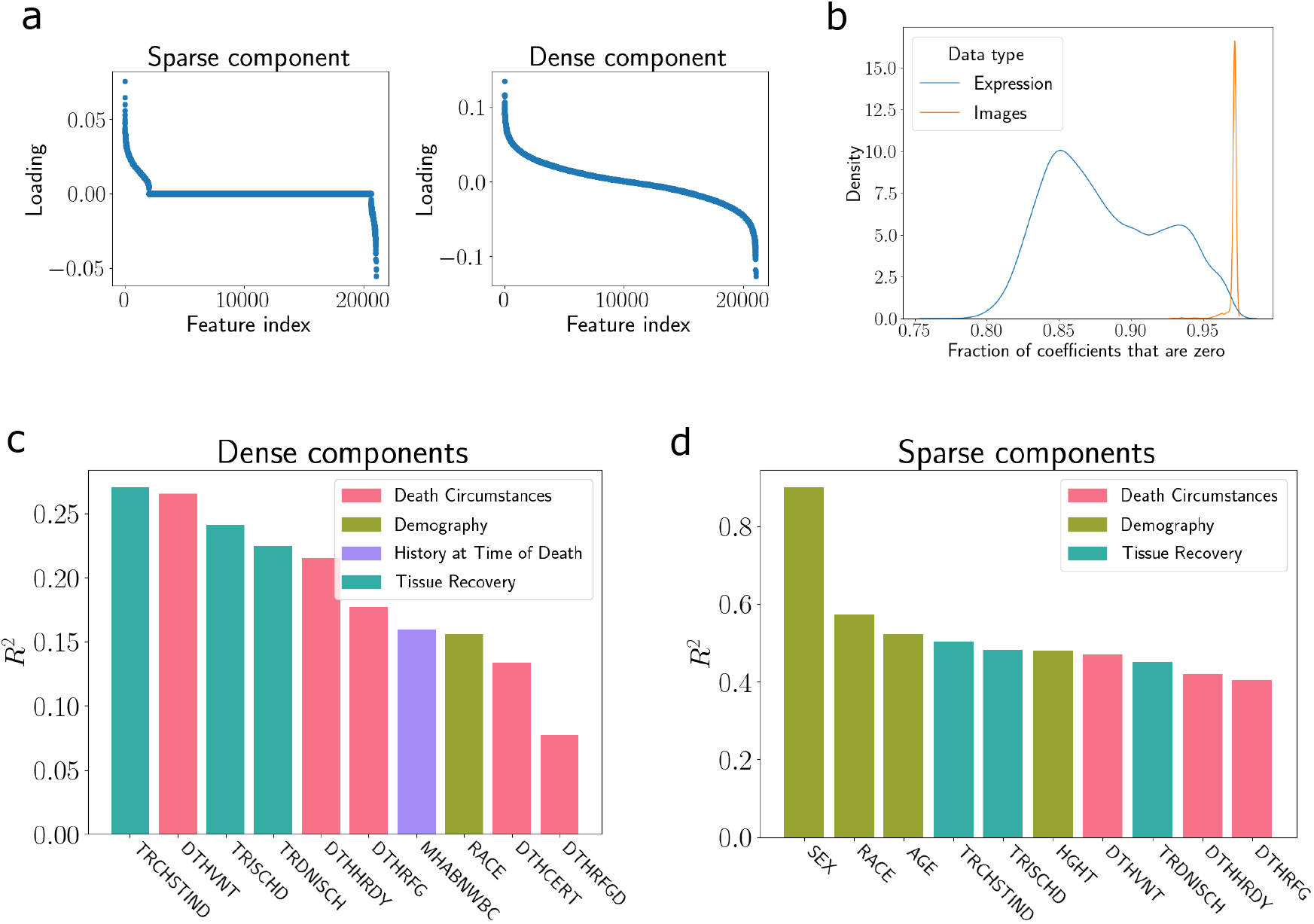
Combinations of sparse and dense components found by SFAmix. **(a)** Loadings values for a sample sparse component (left) and dense component (right). (b) Density plot showing the fraction of zeros in the components corresponding to each feature. Densities are split by whether the feature is gene expression or an image feature. (c) Associations between GTEx metadata and SFAmix dense components. (d) Associations between GTEx metadata and SFAmix sparse components.

To further understand what signal the sparse and dense components capture, we computed the associations between the latent factors and the metadata information. We performed a similar analysis as in Section 2.4, but here we examine the associations separately for sparse and dense components. We find that the dense components tend to be most associated with characteristics related to technical variation and tissue health, such as *time of chest incision* and *ischemic time* (Figure 10c). Meanwhile, the sparse components tend to be highly associated with demographic information such as *sex, race*, and *age* (Figure 10d). These results suggest that the sparse and dense factors — which capture variation in a small number of features and a large number of features, respectively — are encoding complementary sources of variation in these data modalities.

Finally, we investigated whether the SFAmix factors are associated with the genotype of the corresponding samples by performing a QTL analysis. Specifically, we treated the sparse factors as a composite phenotype representing variation in gene expression and histology images. We then ran an analysis similar to our imQTL experiment (Section 2.5). Specifically, we tested for association between each gene variant and latent factor in each tissue using MatrixEQTL (Shabalin, 2012). We found 1, 826 associations across eight tissues (FDR ≤ 0.1). This is substantially more associations than we found in our imQTL analysis — which only tested for associations with image features — suggesting that factors that jointly capture gene expression and histology information could be promising for identifying QTLs.

## 3 Discussion

In this study, we present an analysis of shared variation in gene expression and histology images in a large sample of human tissues from the GTEx v8 project. Our analysis focused on understanding the expression signatures associated with morphological features, as well as the donor and sample characteristics that are related to these signatures.

In general, we found that tissue type is a major driver of variation in both of these modalities, but gene expression contains extensive tissue-specific signal beyond that contained in the image features. However, we also saw substantial variation among image features and gene expression within tissue types. This variation was associated with technical, environmental, and biological factors, including sample ischemic time in liver tissue, donor ventilation status at the time of death in lung tissue, and an image morphology QTL in esophageal mucosa tissue.

While it is evident from our analysis that image morphology and gene expression contain both shared and complementary information, there remain challenges for fully articulating the relationship between them. A crucial future step will be to identify the specific image features associated with various gene expression and genotype features. This type of analysis could be able to identify cell types, tissue structures, and tissue shapes that are related to specific expression phenotypes. We attempted such an analysis in this study by fitting a model to predict pixel-wise expression levels of specific genes within held-out images (Supplement Section 6.3). However, we were unable to achieve consistent accuracy. One solution would be to collect spatially-resolved gene expression data to triangulate the relationship between image morphology and gene expression in order to develop a mapping function between the two data types.

Joint analyses of morphology and molecular features shed light on each modality: morphology can inform us about gene function, and genomic data can inform us about the function of morphological structures. We envision this type of joint analysis being important in future studies of cellular organization and function, and disease.

## 4 Ackowledgments

The authors thank Chuan Gao for help with running SFAmix.

## 5 Methods

### 5.1 Code availability

Code for analyses and data processing is available here: https://github.com/andrewcharlesjones/gtex_image_analysis.

### 5.2 Data

#### 5.2.1 Gene expression

We used the GTEx version 8 (v8) gene expression dataset. Specifically, the transcript per million (TPM) measurements were accessed from the file GTEx Analysis 2017-06-05 v8 RSEMv1.3.0 transcript tpm.gct in the dbGaP public repository. For all analyses, we used the 20, 000 most variable genes. For each gene, we log-transformed, mean-centered, and standardized the data by dividing by its sample standard deviation.

#### 5.2.2 Images

The GTEx v8 histology image dataset is composed of images of tissue slices fixed to slides and stained with hematoxylin and eosin (H&E). Raw images were provided in SVS format. The images used inconsistent magnification, but in general were taken with either 20× apparent magnification (approximately 0.25 microns per pixel) or 40× apparent magnification (approximately 0.50 microns per pixel). The images were imported using the Python interface for the OpenSlide library and then split into 1000 × 1000 pixel tiles. A tile was considered for selection if the mean gray values of itself and the immediately adjacent tiles (above, below, left, and right) were each below (darker than) 180 out of 255. Because of the intractable file size of the full-resolution images, tile selection was actually performed on the 16× lower resolution version of the image, and the region in the full-resolution image corresponding to the selected tile was extracted.

For the convolutional autoencoder analysis, images were further downsampled to create 512 × 512 versions of each image (see below for details).

#### 5.2.3 Sample- and donor-level covariates

The GTEx data contains several metadata related to the samples and the donors. We use 123 donor-level covariates and 37 sample-level covariates.

### 5.3 Metadata variable codes

|  |  |  |  |
| --- | --- | --- | --- |
| *SEX* | Biological sex | *LBHBSAB* | Hepatitis B surface antibody |
| *HGHT* | Height | *DTHRFGD* | Number of hours in refrigeration |
| *TRCHSTIND* | Time of chest incision (minutes) | *TRDNISCH* | Total ischemic time |
| *DTHVNT* | Donor on a ventilator immediately prior to death | *DTHHRDY* | Hardy scale |
| *DTHRFG* | Body refrigerated | *SMRRNART* | rRNA Rate: Ratio of all reads aligned to rRNA regions to total reads |
| *SMRRNANM* | Intronic Rate: The fraction of reads that map within introns | *SMEXNCRT* | Exonic Rate: The fraction of reads that map within exons |
| *SMGNSDTC* | Genes Detected: Total number of genes with at least 5 exon mapping reads | *SMTRSCPT* | Transcripts Detected: Total number of transcripts with at least 5 exon mapping reads |
| *SMSPLTRD* | Split Reads: The number of reads that span an exon-exon boundary | *SMALTALG* | Alternative Alignments: duplicate read entries providing alternative coordinates |
| *SMRIN* | RIN Number | *SMEXPEFF* | Expression Profiling Efficiency: Ratio of exon reads to total reads |
| *LDACC* | Laboratory, Data Analysis and Coordinating Center |  |  |

#### 5.3.1 Human Protein Atlas data

The Human Protein Atlas (HPA) is a Swedish-based research consortium that has generated large-scale profiles of protein activity across many tissues and using many different measurement technologies (Uhlén et al., 2015). We leveraged the “Tissue Atlas” portion of the HPA database, which provides images with spatially localized protein expression estimates across many proteins and human tissues. This is achieved through an immunohistochemical staining process. We downloaded the HPA images using the HPAanalyze R package (Tran et al., 2019), which provides an interface with the HPA online portal.

### 5.4 Image autoencoder

We trained a convolutional autoencoder using the GTEx histology image data using the DCGAN architecture (Radford et al., 2015). We modified the architecture to accept 128×128 images as input. To train the autoencoder, we randomly sampled a 128 × 128 patch of each image on each epoch, and fed a batch of these patches as input. The model was fit in Pytorch using a mean-squared (reconstruction) error and the Adam optimizer (Paszke et al., 2019; Kingma and Ba, 2014).

Once the model was fit, we extracted the activations from the inner-most bottleneck layer for each image — this resulted in a 1024-length vector representation of each image.

### 5.5 Tissue prediction

To assess the extent to which tissue type is encoded in each data modality, we performed a classification analysis with each data type, using a multilayer perceptron with one hidden layer as the classifier (using the Scikit-learn software (Pedregosa et al., 2011)). For gene expression, we used the TPM data as the covariates, and each tissue’s corresponding tissue type as the response. For histology images, we used the autoencoder embeddings as the covariates and the tissue type as the response. We measured the classification performance using the prediction accuracy on a held-out test dataset. We repeated this multiple times through 5-fold cross-validation. We performed a similar analysis to assess the extent to which each CCA method (ImageCCA, PCCA, and DPCCA) captured tissue type in its canonical components. For each method, we extracted the canonical variables (i.e., the latent variable representation) for each sample, and used these as the covariates in the classification analysis.

### 5.6 Ischemic time analysis

- Multiple analyses were performed to assess the extent to which ischemic time is encoded in each dataset.
- First, we performed a regression analysis to assess the overall level of association between each data type and ischemic time in each tissue. Specifically, for each tissue, we ran two linear regression analyses: one in which we regressed ischemic time on the tissue’s gene expression measurements, and another regressing ischemic time onto the tissue’s histology image representation. For the image representation, we used the autoencoder latent representation. We measured the regression performance by computing the Pearson correlation between predicted ischemic time and true ischemic time in a held-out test dataset.
- Second, we performed a more focused analysis on the pancreas tissue data. We first fit a convolutional autoencoder (identical to the one above) on the pancreas data alone. Then, after performing PCA on the latent representations, we correlated each PC with the samples’ ischemic times. Finally, we fit a CCA model using pancreas data alone, and ran a GSEA analysis on the fitted CCA gene coefficients.

### 5.7 ImageCCA

We ran ImageCCA using the code available at https://github.com/daniel-munro/imageCCA, swapping out the ImageCCA autoencoder architecture for the DCGAN128 architecture as described above. ImageCCA uses sparse CCA as implemented by the PMA package in R (Witten et al., 2009). We set the imageCCA hyperparameters as *λ*_*x*_ = 0.1 and *λ*_*y*_ = 0.1, where *λ*_*x*_ and *λ*_*y*_ are the parameters controlling the sparsity penalty for the expression and image data, respectively.

### 5.8 IBFA

IBFA, which is closely related to canonical correlation analysis (CCA), models two datasets **X**^*a*^, **X**^*b*^ with paired samples as being generated from a small set of shared latent variables **z**^*s*^ that explain variation in both, as well as two sets of latent variables **z**^*a*^, **z**^*b*^ that explain variation unique to each dataset. The dimensionalities of the latent variables **z**^*s*^, **z**^*a*^, and **z**^*b*^ are *k*_*s*_, *k*_*a*_, and *k*_*b*_, respectively, where *k*_*s*_, *k*_*a*_, *k*_*b*_ *≪* min(*p, q*). For simplicity, we assume *k*_*s*_ = *k*_*a*_ = *k*_*b*_ in our experiments. Assume that, without loss of generality, **X**^*a*^ and **X**^*b*^ correspond to the image and datasets, respectively. Specifically, if 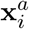 is the *i*th image sample, and 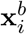 is the corresponding expression sample, PCCA assumes the following generative model for *i* = 1, …, *n*:

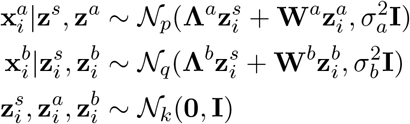

The latent variables 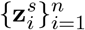 are compressed, low-dimensional representations of each sample that capture information from both the image and expression feature spaces. Ad-ditionally, 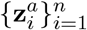 and 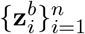 capture low-dimensional variation that is specific to images and expression, respectively. The loadings matrices **Λ**^*a*^ ∈ ℝ^*k×p*^ and **Λ**^*b*^ ∈ ℝ^*k×q*^ capture the linear relationship between the shared latent space and each dataset. Similarly **W**^*a*^ ∈ ℝ^*k×p*^ and **W**^*b*^ ∈ ℝ^*k×q*^ capture the relationship between the view-specific latent spaces and their respective datasets.

**Figure 11.**
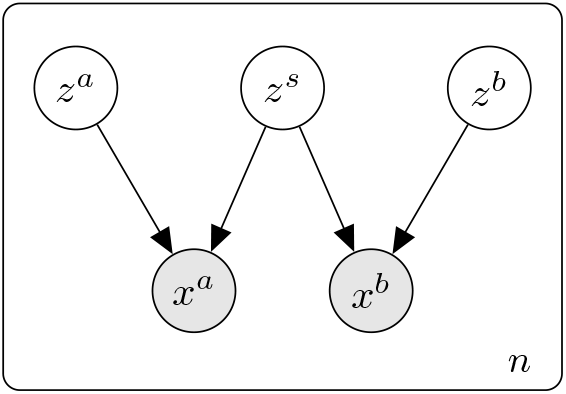
Graphical model for IBFA.

- Probabilistic inter-battery factor analysis (IBFA) (Klami et al., 2013) is an extension of the more well-known probabilistic canonical correlation analysis (PCCA) (Bach and Jordan, 2005). In each of these methods, two datasets with paired samples are represented by a small set of latent variables shared by each. IBFA further includes two extra sets of latent variables that are specific to each dataset.
- The generative model for IBFA is as follows

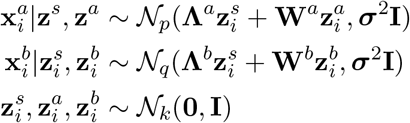

- We perform inference in this model using a mean-field variational approximation. In particular, we approximate the posterior 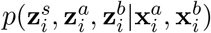 with a fully factorized approximation:

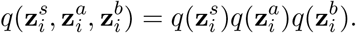

We chose each of the variational distributions to be spherical Gaussians:

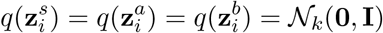

### 5.9 Metadata association analysis

We conducted a series of linear regression analyses to evaluate the association of the joint phenotype with sample- and subject-level covaa matrix of the shared latenrtiates.

Let *n* be the number of samples. For each covariate **y** ∈ ℝ^*n*^, we fit the regression model

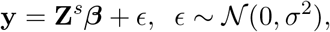

where **Z**^*s*^ ∈ ℝ^*n×k*^ is a matrix of the shared latent variables, and ***β*** ∈ ℝ^*k*^ is a vector of coefficients.

We assessed the strength of the linear regression fit using the *R*^2^ value, which is computed as

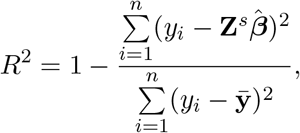

where 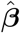 is a vector of the estimated regression coefficients, and 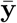 is the mean value of the response variable.

### 5.10 Spatial expression prediction

We trained a convolutional neural network (CNN) to predict the expression value for a given gene at each location in a histology image. Specifically, we used the histology images as input to the CNN, and we used the bulk RNA-seq expression of a given gene as the label. Similar to the autoencoder analysis above, we randomly sampled each image to a 128 × 128 crop on each epoch (but note that all crops had the same bulk gene expression label). We trained the CNN with mean-squared error loss and the Adam optimizer. To predict the expression pattern spatially across a test image, we cropped the image into hundreds of overlapping 128 × 128 tiles, and obtained the CNN’s predicted expression value for each tile. By pasting together these predicted values, we created a heatmap of the spatially-resolved expression predictions for each image.

### 5.11 Gene set enrichment analysis

We performed two types of gene set enrichment analysis (GSEA): a hypergeometric test and a permutation-based test. To run GSEA on the coefficients of ImageCCA — which are sparse and contain only a small number of which are nonzero — we ran a hypergeometric test for each gene set. Specifically, for each gene set, we tested whether the nonzero coefficients were enriched in that set compared to all other sets. We implemented the hypergeometric test using the R package Piano (Väremo et al., 2013). Significance values were corrected using an FDR procedure. To run GSEA on the gene-wise coefficients of PCCA and DPCCA — which are not sparse — we ran a permutation test for each gene set. Specifically, for each gene set, the genes were ranked by their coefficient values, and this ranking was compared to many permuted rankings in order to estimate the enrichment level of that gene set near the top or bottom of the ranked list. We implemented the permutation test using the R package `fgsea` (Korotkevich et al., 2019). We used two classes of gene sets, both provided by MSigDB (Liberzon et al., 2015): the Hallmark gene sets and the GO Biological Process gene sets.

**Supplementary Figure 1.**
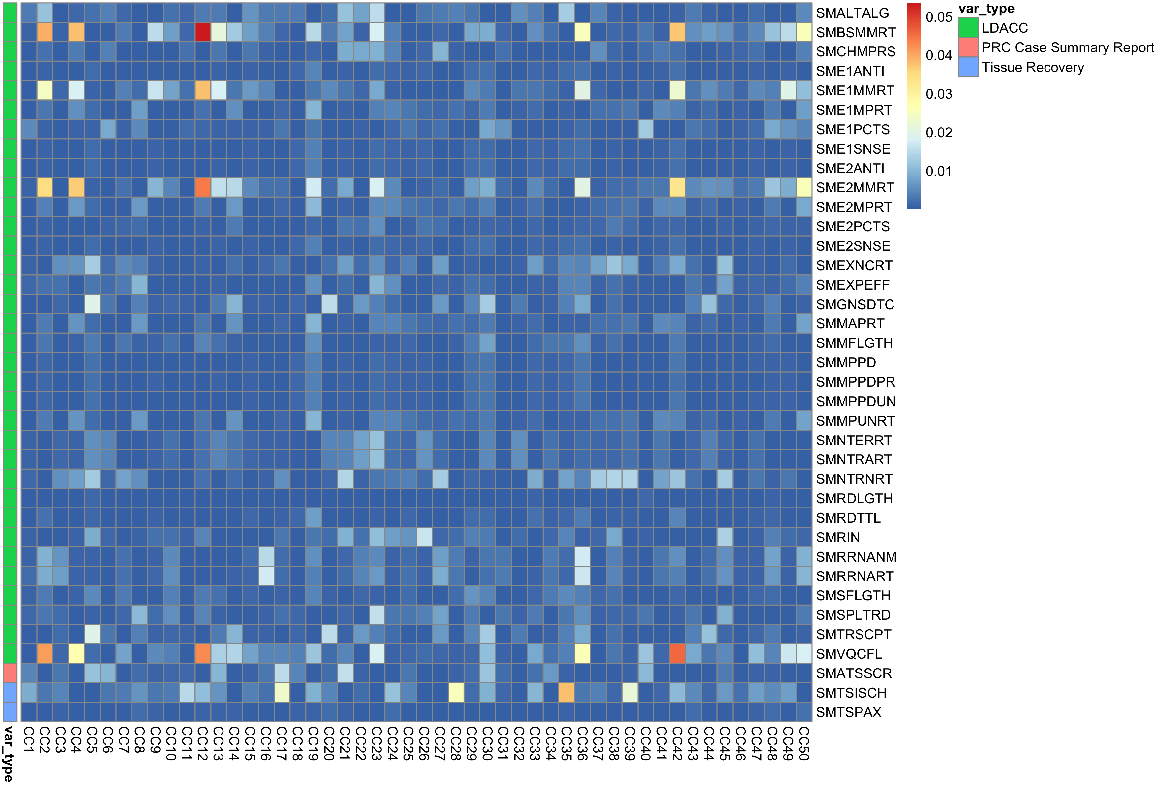
Associations between IBFA latent variables and sample-level meta-data. Each cell represents the *R*^2^ value for each univariate relationship. The color bar on the left represents the category of each metadata variable.

## 6 Supplementary material

### 6.1 Metadata associations

**Supplementary Figure 2.**
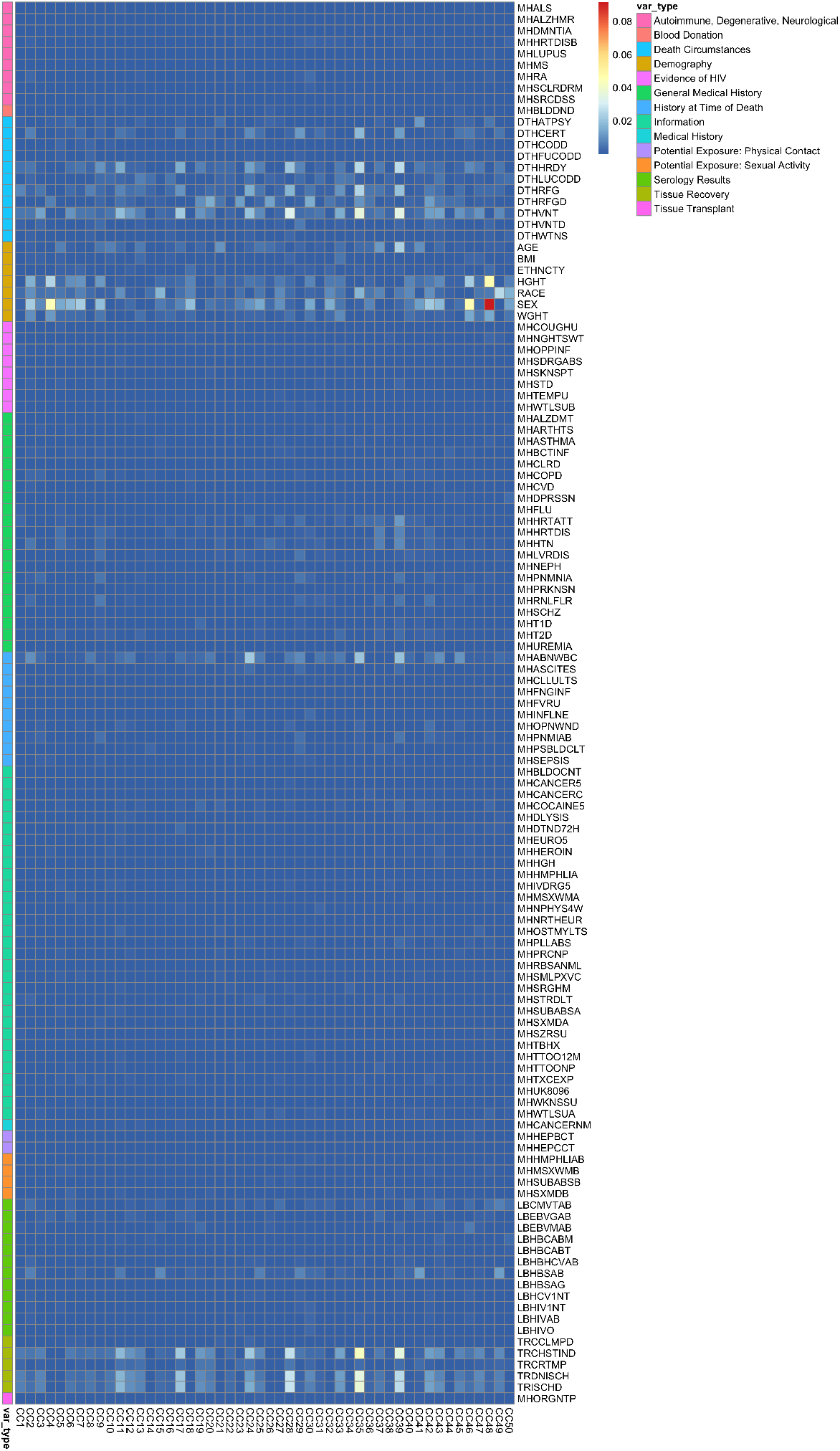
Associations between IBFA latent variables and donor-level meta-data. Each cell represents the *R*^2^ value for each univariate relationship. The color bar on the left represents the category of each metadata variable.

### 6.2 Image QTL hits

| SNP | Image feature | beta | p-value | FDR | tissue | rsid |
| --- | --- | --- | --- | --- | --- | --- |
| chr12_30579577_C_G.b38 | 9 | 0.135 | 3.55e-10 | 0.001 | Brain Cortex | rs34836571 |
| chr12_30579892_A_G.b38 | 9 | 0.135 | 3.55e-10 | 0.001 | Brain Cortex | rs73075778 |
| chr12_30582993_A_G.b38 | 9 | 0.135 | 3.55e-10 | 0.001 | Brain Cortex | rs11614872 |
| chr12_30583690_A_G.b38 | 9 | 0.135 | 3.55e-10 | 0.001 | Brain Cortex | rs73075781 |
| chr12_30586152_T_C.b38 | 9 | 0.135 | 3.55e-10 | 0.001 | Brain Cortex | rs7961681 |
| chr12_30615299_CCAGT_C.b38 | 9 | 0.131 | 3.21e-09 | 0.006 | Brain Cortex | rs35547058 |
| chr12_30626410_C_T.b38 | 9 | 0.138 | 6.61e-09 | 0.006 | Brain Cortex | rs73077865 |
| chr12_30633444_T_C.b38 | 9 | 0.138 | 6.61e-09 | 0.006 | Brain Cortex | rs17486840 |
| chr12_30655998_T_C.b38 | 9 | 0.138 | 6.61e-09 | 0.006 | Brain Cortex | rs55957969 |
| chr12_30656385_C_T.b38 | 9 | 0.138 | 6.61e-09 | 0.006 | Brain Cortex | rs73077870 |
| chr12_30658245_G_T.b38 | 9 | 0.138 | 6.61e-09 | 0.006 | Brain Cortex | rs4931358 |
| chr12_30662569_T_A.b38 | 9 | 0.138 | 6.61e-09 | 0.006 | Brain Cortex | rs73077877 |
| chr12_30676443_C_A.b38 | 9 | 0.138 | 6.61e-09 | 0.006 | Brain Cortex | rs17560141 |
| chr12_30678475_C_G.b38 | 9 | 0.138 | 6.61e-09 | 0.006 | Brain Cortex | rs73077884 |
| chr12_30683436_T_C.b38 | 9 | 0.138 | 6.61e-09 | 0.006 | Brain Cortex | rs73077887 |
| chr12_30689456_C_T.b38 | 9 | 0.138 | 6.61e-09 | 0.006 | Brain Cortex | rs11608803 |
| chr12_30693773_C_T.b38 | 9 | 0.138 | 6.61e-09 | 0.006 | Brain Cortex | rs73077893 |
| chr12_30614207_G_C.b38 | 9 | 0.127 | 7.06e-09 | 0.006 | Brain Cortex | rs3850959 |
| chr11_67518604_T_C.b38 | 1 | 0.166 | 3.02e-09 | 0.009 | Skin Sun Exposed Lower leg | rs682208 |
| chr2_196254354_T_G.b38 | 1 | 0.117 | 4.86e-10 | 0.021 | Skin Sun Exposed Lower leg | rs4372925 |
| chr2_196205718_G_T.b38 | 1 | -0.115 | 1.48e-09 | 0.03 | Skin Sun Exposed Lower leg | rs1455815 |
| chr2_196252433_G_C.b38 | 1 | 0.11 | 3.02e-09 | 0.03 | Skin Sun Exposed Lower leg | rs4850689 |
| chr2_196253298_T_C.b38 | 1 | 0.11 | 3.02e-09 | 0.03 | Skin Sun Exposed Lower leg | rs1869796 |
| chr2_196216491_T_C.b38 | 1 | -0.113 | 3.47e-09 | 0.03 | Skin Sun Exposed Lower leg | rs6707519 |
| chr2_196250030_A_G.b38 | 1 | 0.107 | 6.64e-09 | 0.049 | Skin Sun Exposed Lower leg | rs4850687 |
| chr2_196254354_T_G.b38 | 1 | 0.117 | 4.86e-10 | 0.013 | Skin Sun Exposed Lower leg | rs4372925 |
| chr2_196205718_G_T.b38 | 1 | -0.115 | 1.48e-09 | 0.019 | Skin Sun Exposed Lower leg | rs1455815 |
| chr2_196252433_G_C.b38 | 1 | 0.11 | 3.02e-09 | 0.019 | Skin Sun Exposed Lower leg | rs4850689 |
| chr2_196253298_T_C.b38 | 1 | 0.11 | 3.02e-09 | 0.019 | Skin Sun Exposed Lower leg | rs1869796 |
| chr2_196216491_T_C.b38 | 1 | -0.113 | 3.47e-09 | 0.019 | Skin Sun Exposed Lower leg | rs6707519 |
| chr2_196250030_A_G.b38 | 1 | 0.107 | 6.64e-09 | 0.03 | Skin Sun Exposed Lower leg | rs4850687 |
| chr2_196254354_T_G.b38 | 1 | 0.117 | 4.86e-10 | 0.02 | Skin Sun Exposed Lower leg | rs4372925 |
| chr2_196205718_G_T.b38 | 1 | -0.115 | 1.48e-09 | 0.028 | Skin Sun Exposed Lower leg | rs1455815 |
| chr2_196252433_G_C.b38 | 1 | 0.11 | 3.02e-09 | 0.028 | Skin Sun Exposed Lower leg | rs4850689 |
| chr2_196253298_T_C.b38 | 1 | 0.11 | 3.02e-09 | 0.028 | Skin Sun Exposed Lower leg | rs1869796 |
| chr2_196216491_T_C_b38 | 1 | -0.113 | 3.47e-09 | 0.028 | Skin Sun Exposed Lower leg | rs6707519 |
| chr2_196250030_A_G_b38 | 1 | 0.107 | 6.64e-09 | 0.045 | Skin Sun Exposed Lower leg | rs4850687 |
| chr2_196254354_T_G_b38 | 1 | 0.117 | 4.86e-10 | 0.029 | Skin Sun Exposed Lower leg | rs4372925 |
| chr2_196205718_G_T_b38 | 1 | -0.115 | 1.48e-09 | 0.041 | Skin Sun Exposed Lower leg | rs1455815 |
| chr2_196252433_G_C_b38 | 1 | 0.11 | 3.02e-09 | 0.041 | Skin Sun Exposed Lower leg | rs4850689 |
| chr2_196253298_T_C_b38 | 1 | 0.11 | 3.02e-09 | 0.041 | Skin Sun Exposed Lower leg | rs1869796 |
| chr2_196216491_T_C_b38 | 1 | -0.113 | 3.47e-09 | 0.041 | Skin Sun Exposed Lower leg | rs6707519 |
| chr2_196250030_A_G_b38 | 1 | 0.107 | 6.64e-09 | 0.065 | Skin Sun Exposed Lower leg | rs4850687 |
| chr19_3766809_C_T_b38 | 919 | -0.001 | 5.26e-09 | 0.078 | Stomach | rs55927187 |
| chr19_3766579_G_A_b38 | 919 | -0.001 | 9.19e-09 | 0.078 | Stomach | rs56363913 |
| chr19_3766809_C_T_b38 | 919 | -0.001 | 5.26e-09 | 0.113 | Stomach | rs55927187 |
| chr19_3766579_G_A_b38 | 919 | -0.001 | 9.19e-09 | 0.113 | Stomach | rs56363913 |
| chr9_37749476_C_CAG_b38 | 661 | 0.007 | 8.68e-09 | 0.196 | Thyroid | rs60084650 |
| chr9_37749476_C_CAG_b38 | 661 | 0.007 | 8.68e-09 | 0.397 | Thyroid | rs60084650 |
| chr9_37749476_C_CAG_b38 | 661 | 0.007 | 8.68e-09 | 0.546 | Thyroid | rs60084650 |
| chr9_37749476_C_CAG_b38 | 661 | 0.007 | 8.68e-09 | 0.059 | Thyroid | rs60084650 |
| chr1_48587261_T_C_b38 | 280 | 0.008 | 8.24e-09 | 0.283 | Muscle Skeletal | rs2803277 |
| chr4_7967867_G_A_b38 | 1007 | 0.0 | 4.58e-09 | 0.092 | Esophagus Mucosa | rs10008860 |

[utbl]

### 6.3 “Painting” expression onto images

Given that we found several clear relationships between gene expression and morphological features, we asked whether we could infer a dense spatial labeling of gene expression on a tissue sample from histology images alone. In other words, given an image of a tissue sample without the corresponding gene expression measurement, can we infer the spatial layout of the expression of a gene across the tissue slice?

#### 6.3.1 Dense spatial labeling of gene expression

To test this, we fit a convolutional neural network to predict the bulk expression measurement for each gene from the corresponding histology image. Specifically, we split up each image into smaller, overlapping tiles, and trained the CNN to predict the corresponding expression value from these tiles. Then, passing the tiles of a test image through the fitted CNN provides a dense predicted label of a gene’s expression values across the image.

Given that our expression measurements were bulk RNA-seq samples that were did not contain any spatial information, it is impossible to verify our predictions with this data alone, and we thus show our experiments as a proof-of-concept. Nevertheless, we found that the CNN was able to identify large morphological structures and map these onto expression patterns. For example, in Thyroid tissue, we found the predicted location of expression of *FOXE1* to correlate with the location of cells (Supplementary Figure 3).

**Supplementary Figure 3:**
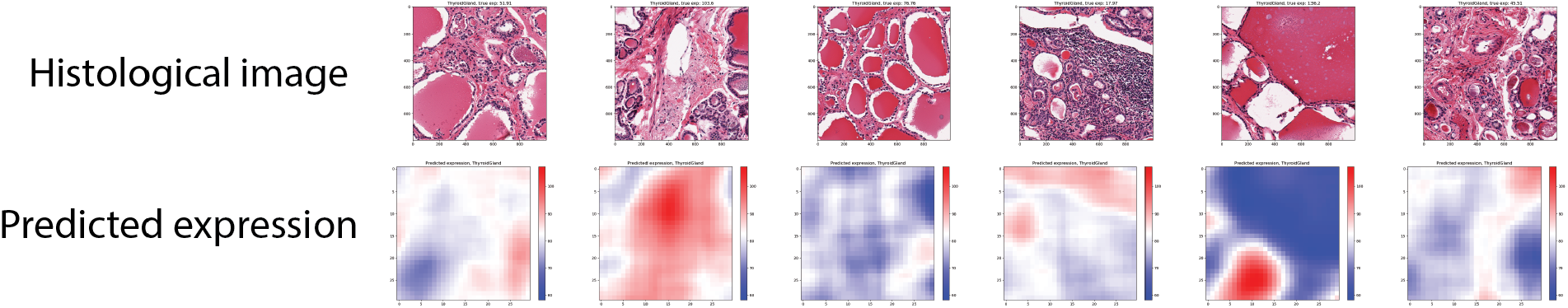
Predictions from a CNN trained to localize expression patterns within individual histology images.

#### 6.3.2 Validating with Human Protein Atlas data

To attempt to validate this proof-of-concept, we tested the CNN on a dataset with spatially labeled expression. In particular, we leveraged data from the Human Protein Atlas (HPA) (Uhlén et al., 2015), which contains measurements of spatially localized protein expression. For this test, we again fit the CNN on thyroid data from GTEx for the gene *INHA*. Here, we used grayscale images in order to account for color differences between the GTEx images and HPA images. We then computed spatially localized predictions for a sample from the HPA.

We find that the predictions generally resemble the ground truth of protein expression (Supplementary Figure 4). We suspect that recent advancements in spatial gene expression profiling could further enhance this analysis (Ståhl et al., 2016; Rodriques et al., 2019).

**Supplementary Figure 4.**
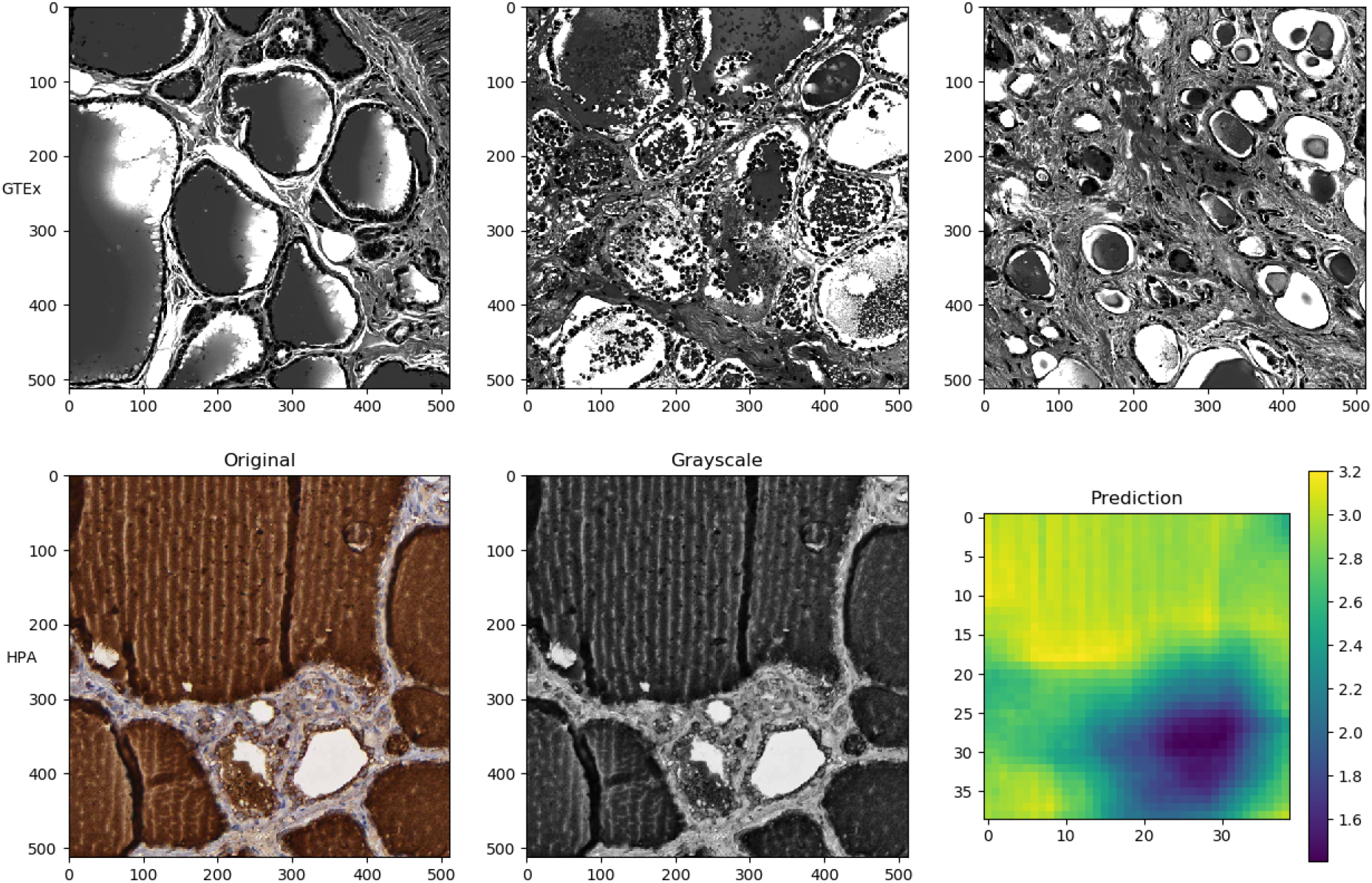
Predicting *INHA* expression in thyroid tissues spatially using the Human Protein Atlas data.

## References

[1] Frank W Albert and Leonid Kruglyak. The role of regulatory variation in complex traits and disease. Nature Reviews Genetics, 16(4):197–212, 2015.

[2] Teresa Araújo, Guilherme Aresta, Eduardo Castro, José Rouco, Paulo Aguiar, Catarina Eloy, António Polónia, and Aurélio Campilho. Classification of breast cancer histology images using convolutional neural networks. PloS One, 12(6):e0177544, 2017.

[3] Guilherme Aresta, Teresa Araújo, Scotty Kwok, Sai Saketh Chennamsetty, Mohammed Safwan, Varghese Alex, Bahram Marami, Marcel Prastawa, Monica Chan, Michael Donovan, et al. Bach: Grand challenge on breast cancer histology images. Medical Image Analysis, 56:122–139, 2019.

[4] Jordan T Ash, Gregory Darnell, Daniel Munro, and Barbara E Engelhardt. Joint analysis of expression levels and histological images identifies genes associated with tissue morphology. Nature Communications, 12(1):1–12, 2021.

[5] Francis R Bach and Michael I Jordan. A probabilistic interpretation of canonical correlation analysis. Technical Report, 2005.

[6] Laura Barisoni, Kyle J Lafata, Stephen M Hewitt, Anant Madabhushi, and Ulysses GJ Balis. Digital pathology and computational image analysis in nephropathology. Nature Reviews Nephrology, 16(11):669–685, 2020.

[7] Joseph D Barry, Maud Fagny, Joseph N Paulson, Hugo JWL Aerts, John Platig, and John Quackenbush. Histopathological image QTL discovery of immune infiltration variants. iScience, 5:80–89, 2018.

[8] Andrew H Beck, Ankur R Sangoi, Samuel Leung, Robert J Marinelli, Torsten O Nielsen, Marc J Van De Vijver, Robert B West, Matt Van De Rijn, and Daphne Koller. Systematic analysis of breast cancer morphology uncovers stromal features associated with survival. Science Translational Medicine, 3(108):108ra113–108ra113, 2011.

[9] Babak Ehteshami Bejnordi, Mitko Veta, Paul Johannes Van Diest, Bram Van Ginneken, Nico Karssemeijer, Geert Litjens, Jeroen AWM Van Der Laak, Meyke Hermsen, Quirine F Manson, Maschenka Balkenhol, et al. Diagnostic assessment of deep learning algorithms for detection of lymph node metastases in women with breast cancer. Journal of the American Medical Association, 318(22):2199–2210, 2017.

[10] Latarsha J Carithers, Kristin Ardlie, Mary Barcus, Philip A Branton, Angela Britton, Stephen A Buia, Carolyn C Compton, David S DeLuca, Joanne Peter-Demchok, Ellen T Gelfand, et al. A novel approach to high-quality postmortem tissue procurement: The GTEx project. Biopreservation and Biobanking, 13(5):311–319, 2015.

[11] Dan C Cireşan, Alessandro Giusti, Luca M Gambardella, and Jürgen Schmidhuber. Mitosis detection in breast cancer histology images with deep neural networks. In International conference on medical image computing and computer-assisted intervention, pages 411–418. Springer, 2013.

[12] Ana Conesa, Pedro Madrigal, Sonia Tarazona, David Gomez-Cabrero, Alejandra Cervera, Andrew McPherson, Michal Wojciech Szczésniak, Daniel J Gaffney, Laura L Elo, Xuegong Zhang, et al. A survey of best practices for RNA-seq data analysis. Genome Biology, 17 (1):1–19, 2016.

[13] William Cookson, Liming Liang, Gonçalo Abecasis, Miriam Moffatt, and Mark Lathrop. Mapping complex disease traits with global gene expression. Nature Reviews Genetics, 10 (3):184–194, 2009.

[14] Lee AD Cooper, Jun Kong, David A Gutman, Fusheng Wang, Jingjing Gao, Christina Appin, Sharath Cholleti, Tony Pan, Ashish Sharma, Lisa Scarpace, et al. Integrated morphologic analysis for the identification and characterization of disease subtypes. Journal of the American Medical Informatics Association, 19(2):317–323, 2012.

[15] Yu Fu, Alexander W Jung, Ramon Viñas Torne, Santiago Gonzalez, Harald Vöhringer, Artem Shmatko, Lucy R Yates, Mercedes Jimenez-Linan, Luiza Moore, and Moritz Gerstung. Pan-cancer computational histopathology reveals mutations, tumor composition anMetadata variable codesd prognosis. Nature Cancer, 1(8):800–810, 2020.

[16] Thomas J Fuchs and Joachim M Buhmann. Computational pathology: Challenges and promises for tissue analysis. Computerized Medical Imaging and Graphics, 35(7-8):515–530, 2011.

[17] Chuan Gao, Christopher D Brown, and Barbara E Engelhardt. A latent factor model with a mixture of sparse and dense factors to model gene expression data with confounding effects. arXiv preprint 1310.4792, 2013.

[18] Daniel Gratz, Alexander J Winkle, Alyssa Dalic, Sathya D Unudurthi, and Thomas J Hund. Computational tools for automated histological image analysis and quantification in cardiac tissue. MethodsX, 7:100755, 2020.

[19] The GTEx Consortium et al. Genetic effects on gene expression across human tissues. Nature, 550(7675):204–213, 2017.

[20] The GTEx Consortium et al. The GTEx consortium atlas of genetic regulatory effects across human tissues. Science, 369(6509):1318–1330, 2020.

[21] Gregory Gundersen, Bianca Dumitrascu, Jordan T Ash, and Barbara E Engelhardt. End-toend training of deep probabilistic CCA on paired biomedical observations. In Uncertainty in Artificial Intelligence, 2019.

[22] Kaiming He, Xiangyu Zhang, Shaoqing Ren, and Jian Sun. Deep residual learning for image recognition. In Proceedings of the IEEE conference on computer vision and pattern recognition, pages 770–778, 2016.

[23] Friederike Hoellen, Athina Kostara, Thomas Karn, Uwe Holtrich, Ahmed El-Balat, Mike Otto, Achim Rody, and Lars C Hanker. Trefoil factor 3 expression in epithelial ovarian cancer exerts a minor effect on clinicopathological parameters. Molecular and Clinical Oncology, 5(4):422–428, 2016.

[24] Harold Hotelling. Relations between two sets of variates. In Breakthroughs in statistics, pages 162–190. Springer, 1992.

[25] The ICGC/TCGA Pan-Cancer Analysis of Whole Genomes Consortium et al. Pan-cancer analysis of whole genomes. Nature, 578(7793):82, 2020.

[26] Diederik P Kingma and Jimmy Ba. Adam: A method for stochastic optimization. arXiv preprint 1412.6980, 2014.

[27] Arto Klami, Seppo Virtanen, and Samuel Kaski. Bayesian canonical correlation analysis. Journal of Machine Learning Research, 14(Apr):965–1003, 2013.

[28] Daisuke Komura and Shumpei Ishikawa. Machine learning methods for histopathological image analysis. Computational and Structural Biotechnology Journal, 16:34–42, 2018.

[29] Gennady Korotkevich, Vladimir Sukhov, and Alexey Sergushichev. Fast gene set enrichment analysis. BioRxiv, page 060012, 2019.

[30] Sonal Kothari, John H Phan, Todd H Stokes, and May D Wang. Pathology imaging informatics for quantitative analysis of whole-slide images. Journal of the American Medical Informatics Association, 20(6):1099–1108, 2013.

[31] Alex Krizhevsky, Ilya Sutskever, and Geoffrey E Hinton. Imagenet classification with deep convolutional neural networks. Advances in Neural Information Processing Systems, 25:1097–1105, 2012.

[32] Alex Krizhevsky, Ilya Sutskever, and Geoffrey E Hinton. Imagenet classification with deep convolutional neural networks. Communications of the ACM, 60(6):84–90, 2017.

[33] Arthur Liberzon, Chet Birger, Helga Thorvaldsdóttir, Mahmoud Ghandi, Jill P Mesirov, and Pablo Tamayo. The molecular signatures database hallmark gene set collection. Cell systems, 1(6):417–425, 2015.

[34] Li Liu, Xiangshun Li, Rui Yuan, Honghong Zhang, Lixia Qiang, Jingling Shen, and Shoude Jin. Associations of ABHD2 genetic variations with risks for chronic obstructive pulmonary disease in a Chinese Han population. PloS One, 10(4):e0123929, 2015.

[35] Jonathan Masci, Ueli Meier, Dan Cireşan, and Jürgen Schmidhuber. Stacked convolutional auto-encoders for hierarchical feature extraction. In International Conference on Artificial Neural Networks, pages 52–59. Springer, 2011.

[36] Leland McInnes, John Healy, and James Melville. UMAP: Uniform manifold approximation and projection for dimension reduction. arXiv preprint 1802.03426, 2018.

[37] Erick Moen, Dylan Bannon, Takamasa Kudo, William Graf, Markus Covert, and David Van Valen. Deep learning for cellular image analysis. Nature Methods, pages 1–14, 2019.

[38] Alexandr Nedzved, Sergey Ablameyko, and Ioannis Pitas. Morphological segmentation of histology cell images. In Proceedings 15th International Conference on Pattern Recognition. ICPR-2000, volume 1, pages 500–503. IEEE, 2000.

[39] Cancer Genome Atlas Network et al. Comprehensive molecular portraits of human breast tumours. Nature, 490(7418):61, 2012.

[40] Adam Paszke, Sam Gross, Francisco Massa, Adam Lerer, James Bradbury, Gregory Chanan, Trevor Killeen, Zeming Lin, Natalia Gimelshein, Luca Antiga, et al. PyTorch: An imperative style, high-performance deep learning library. In Advances in Neural Information Processing Systems, pages 8026–8037, 2019.

[41] F. Pedregosa, G. Varoquaux, A. Gramfort, V. Michel, B. Thirion, O. Grisel, M. Blondel, P. Prettenhofer, R. Weiss, V. Dubourg, J. Vanderplas, A. Passos, D. Cournapeau, M. Brucher, M. Perrot, and E. Duchesnay. Scikit-learn: Machine learning in Python. Journal of Machine Learning Research, 12:2825–2830, 2011.

[42] Hady Ahmady Phoulady, Dmitry B Goldgof, Lawrence O Hall, and Peter R Mouton. Nucleus segmentation in histology images with hierarchical multilevel thresholding. In Medical Imaging 2016: Digital Pathology, volume 9791, page 979111. International Society for Optics and Photonics, 2016.

[43] Alec Radford, Luke Metz, and Soumith Chintala. Unsupervised representation learning with deep convolutional generative adversarial networks. arXiv preprint 1511.06434, 2015.

[44] Matthew V Rockman and Leonid Kruglyak. Genetics of global gene expression. Nature Reviews Genetics, 7(11):862–872, 2006.

[45] Samuel G Rodriques, Robert R Stickels, Aleksandrina Goeva, Carly A Martin, Evan Murray, Charles R Vanderburg, Joshua Welch, Linlin M Chen, Fei Chen, and Evan Z Macosko. Slide-seq: A scalable technology for measuring genome-wide expression at high spatial resolution. Science, 363(6434):1463–1467, 2019.

[46] Michael L Salmans, Fang Zhao, and Bogi Andersen. The estrogen-regulated anterior gradient 2 (AGR2) protein in breast cancer: a potential drug target and biomarker. Breast Cancer Research, 15(2):1–14, 2013.

[47] Andreas Schroeder, Odilo Mueller, Susanne Stocker, Ruediger Salowsky, Michael Leiber, Marcus Gassmann, Samar Lightfoot, Wolfram Menzel, Martin Granzow, and Thomas Ragg. The RIN: an RNA integrity number for assigning integrity values to RNA measurements. BMC Molecular Biology, 7(1):1–14, 2006.

[48] Andrey A Shabalin. Matrix eQTL: ultra fast eQTL analysis via large matrix operations. Bioinformatics, 28(10):1353–1358, 2012.

[49] Sudhir Sornapudi, Ronald Joe Stanley, William V Stoecker, Haidar Almubarak, Rodney Long, Sameer Antani, George Thoma, Rosemary Zuna, and Shelliane R Frazier. Deep learning nuclei detection in digitized histology images by superpixels. Journal of Pathology Informatics, 9, 2018.

[50] Patrik L Ståhl, Fredrik Salmén, Sanja Vickovic, Anna Lundmark, José Fernández Navarro, Jens Magnusson, Stefania Giacomello, Michaela Asp, Jakub O Westholm, Mikael Huss, et al. Visualization and analysis of gene expression in tissue sections by spatial transcriptomics. Science, 353(6294):78–82, 2016.

[51] Vaishnavi Subramanian, Benjamin Chidester, Jian Ma, and Minh N Do. Correlating cellular features with gene expression using CCA. In 2018 IEEE 15th International Symposium on Biomedical Imaging (ISBI 2018), pages 805–808. IEEE, 2018.

[52] Yan Sun, Jiamin Yang, Shujing Zhang, Yushan Gao, Yixi Jin, Min Wang, Lei Ni, Zhenyi Wang, Pengfei Kang, Yan Qiu, et al. Effect of mechanical ventilation on urine volume and expression of aquaporins in rabbits. Journal of Traditional Chinese Medical Sciences, 4(3):272–279, 2017.

[53] Anh Nhat Tran, Alex M Dussaq, Timothy Kennell, Christopher D Willey, and Anita B Hjelmeland.HPAanalyze: an R package that facilitates the retrieval and analysis of the Human Protein Atlas data. BMC Bioinformatics, 20(1):463, 2019.

[54] Mathias Uhlén, Linn Fagerberg, Björn M Hallström, Cecilia Lindskog, Per Oksvold, Adil Mardinoglu, Åsa Sivertsson, Caroline Kampf, Evelina Sjöstedt, Anna Asplund, et al. Tissue-based map of the human proteome. Science, 347(6220), 2015.

[55] David A Van Valen, Takamasa Kudo, Keara M Lane, Derek N Macklin, Nicolas T Quach, Mialy M DeFelice, Inbal Maayan, Yu Tanouchi, Euan A Ashley, and Markus W Covert. Deep learning automates the quantitative analysis of individual cells in live-cell imaging experiments. PLoS Computational Biology, 12(11):e1005177, 2016.

[56] Leif Väremo, Jens Nielsen, and Intawat Nookaew. Enriching the gene set analysis of genome-wide data by incorporating directionality of gene expression and combining statistical hypotheses and methods. Nucleic Acids Research, 41(8):4378–4391, 2013.

[57] John N Weinstein, Eric A Collisson, Gordon B Mills, Kenna R Mills Shaw, Brad A Ozenberger, Kyle Ellrott, Ilya Shmulevich, Chris Sander, Joshua M Stuart, Cancer Genome Atlas Research Network, et al. The cancer genome atlas pan-cancer analysis project. Nature Genetics, 45(10):1113, 2013.

[58] Daniela M Witten, Robert Tibshirani, and Trevor Hastie. A penalized matrix decomposition, with applications to sparse principal components and canonical correlation analysis. Biostatistics, 10(3):515–534, 2009.

[59] Wenjing Wu, Tushar D Bhagat, Xue Yang, Jee Hoon Song, Yulan Cheng, Rachana Agarwal, John M Abraham, Sariat Ibrahim, Matthias Bartenstein, Zulfiqar Hussain, et al. Hypomethylation of noncoding DNA regions and overexpression of the long noncoding RNA, AFAP1-AS1, in Barrett’s esophagus and esophageal adenocarcinoma. Gastroenterology, 144(5):956–966, 2013.

